# Cortical facilitation of somatosensory inputs using gravity-related tactile information in patients with bilateral vestibular loss

**DOI:** 10.1101/2022.04.19.488449

**Authors:** Marie Fabre, Laura Beullier, Chloé Sutter, Amirezza Krebritchi, Pascale Chavet, Martin Simoneau, Michel Toupet, Jean Blouin, Laurence Mouchnino

**Affiliations:** Aix Marseille Univ. CNRS, Laboratoire de Neurosciences Cognitives, FR 3C, Marseille, France; Centre interdisciplinaire de recherche en réadaptation et intégration sociale (CIRRIS) du CIUSSS de la Capitale Nationale, Québec, QC G1M 2S8, Canada; Département de kinésiologie, Faculté de médecine, Université Laval, Québec, QC G1V 0A6, Canada; Aix Marseille Univ, CNRS, Institut des Sciences du Mouvement, Marseille, France; IRON, Institut de Recherche en Oto-Neurologie, Paris, France; Centre d’Explorations Fonctionnelles Oto-Neurologiques, Paris, France

**Keywords:** Balance, Plantar sole afferents, bilateral vestibular loss, tactile compensation, EEG, SEP, time frequency analysis

## Abstract

A few years after their bilateral vestibular loss, individuals usually show a motor repertoire that is almost back to normal. This recovery is thought to involve an up-regulation of the visual and proprioceptive information that compensates for the lack of vestibular information. Here, we investigated whether plantar tactile inputs, which provide body information relative to the ground and to the Earth-vertical, contribute to this compensation. More specifically, we tested the hypothesis that somatosensory cortex response to electric stimulation of the plantar sole in standing adults will be greater in patients (n = 10) with bilateral vestibular loss than in an aged-matched healthy group (n = 10). Showing significant greater somatosensory evoked potentials (i.e., P_1_N_1_) in patients than in controls, the electroencephalographic recordings supported this hypothesis. Furthermore, we found evidence that increasing the differential pressure between both feet, by adding a 1 kg mass at each pending wrist, enhanced the internal representation of body orientation and motion relative to a gravitational reference frame. The large decreased in alpha/beta power in the right posterior parietal cortex (and not in the left) is in line with this assumption. Finally, our behavioral analyses showed smaller body sway oscillations for patients, likely originated from a tactile-based control strategy. Conversely, healthy subjects showed smaller head oscillations suggesting a vestibular-based control strategy, the head serving as a reference for balance control.

**Highlights:**

- Somatosensory cortex excitability is greater in patients with bilateral vestibular loss than in aged-matched healthy individuals
- To control balance, healthy individuals “locked” the head while vestibular patients “locked” their pelvis
- For vestibular patients, increasing loading/unloading mechanism enhances the internal representation of body state in the posterior parietal cortex

## Introduction

When standing upright, balance can be jeopardized by gravitational force if not properly controlled. This control is thought to involve feedforward processes, and to rely on the use and the quality of internal representations of body orientation and motion in space (Borel et al., 2008; Massion, 1992; Pfeiffer et al., 2014 for reviews). Observations in patients with major vestibular hypofunction (VH) suggest that the gravitational reference frame is crucial for encoding such body-related information. Investigating balance control in VH provides a unique opportunity to assess the role of this geocentric frame of reference for balance control. The vestibular system is paramount for constructing the geocentric frame of reference as it senses linear and angular body accelerations (including Earth gravitational force). The alteration of this reference frame would be responsible for the trunk deviation relative to the earth-vertical that is commonly observed when VH individuals stand on a moving surface (Horak et al., 2002). Similar body deviations were also reported in cats with bilateral labyrinthectomy during comparable stance perturbations (Macpherson et al., 2007). With impaired otoliths-based gravitational reference (see Day & Fitzpatrick, 2005 for a review), the nervous system would be able to detect body motion relative to the supporting surface (through the processing of somatosensory inputs) but would be impaired when determining how the body is moving relative to the earth-vertical (see Horak et al., 2002). Other cat studies found delayed postural responses following translations of the supporting surface when somatosensory inputs were removed (with a large dose of Pyridoxine, Stapley et al., 2002), but not after bilateral labyrinthectomy (Inglis & Macpherson, 1995). Therefore, the greater incidence of falls and the deteriorated quality of life in individuals with VH (Herdman et al., 2000; Ward et al., 2013; Zingler et al., 2008; Pavlou et al., 2006) even years after the onset of their first symptoms, cannot solely be attributed to a deficit in postural reactions. Rather, vestibular disorder complications could be linked to an impaired representation of body orientation and motion relative to the external world (i.e., earth gravity vertical).

It is well known that sensory loss or decrease in the reliability of sensory information can alter the processing of the spared sensory modalities. In such cases, intact sensory inputs are upregulated, and are given more weight by the central nervous system (Angelaki & Laurens, 2020; Bronstein, 2016; Lopez et al., 2007; Toupet et al., 2017). Surprisingly, despite the fact that the plantar mechanoreceptors provide useful information about body position and velocity relative to the ground and to the earth-vertical (e.g., Carriot et al., 2004; Morasso & Schiepatti, 1999; Mouchnino & Blouin, 2013), it has yet to be determined whether cutaneous inputs are enhanced in absence of vestibular information. Upregulation of cutaneous inputs in VH individuals would support studies showing that the brain can selectively increase the functional gain of task-relevant sensory inputs, including those from somatosensory systems during challenging equilibrium tasks (Duysens et al., 1995; Mouchnino et al., 2015; Saradjian et al., 2013; 2019).

The primary aim of the present study was to explore whether the response of the somatosensory cortex to foot cutaneous stimulation, increases in individuals with bilateral vestibular hypofunction. We tested this hypothesis by measuring the amplitude of the P_1_N_1_ cortical response, extracted from the electroencephalogram, following the electric stimulation of the plantar sole (i.e., somatosensory-evoked potential (SEP) technique). We reasoned that the amplitude of the P_1_N_1_ should be a key variable in comparing the weighting of foot cutaneous inputs between VH and healthy control individuals.

Importantly, whether or not foot cutaneous inputs were to be upregulated in individuals with VH, balance-related issues are still reported in patients with vestibular hypofunction (Herdman et al., 2000; Ward et al., 2013; Zingler et al., 2008). This suggests that the sensory compensation if present, would not suffice to create a reliable gravitational reference frame. Particularly, the mechanical action of gravity exerted on all body parts ultimately generates forces on the supporting surface depending on body configuration (Winter et al., 1996). Among these forces, the shear forces that are small relative to the weight force during natural quiet standing, are readily detectable by the tactile receptors (Morasso & Schieppati, 1999; Knellwolf et al., 2018). Along these lines, our second aim was to test if increasing the differential forces between both feet would enhance the internal representation of body orientation and motion relative to a gravitational reference frame in individuals with VH. During upright standing, the pressure exerted under the loading foot increases while the pressure exerted under the unloading foot simultaneously decreases (Winter et al., 1996). This loading/unloading mechanism provides gravity-related tactile information as slow adapting afferent fibers (e.g., Merkel and Ruffini) from the plantar sole increase their firing rate resulting from changes in normal force applied to the skin (Vallbo & Johansson, 1984; see Macefield, 2005 for a review). Therefore, the impulse discharge pattern alternates between the feet depending on the loading or unloading. This phenomenon is reminiscent of the effect of body tilts on vestibular neurons in healthy individuals. The neurons located in the labyrinth ipsilateral to the tilt increase their discharge rates while neurons of the contralateral labyrinth concomitantly decrease theirs (Fernandez & Goldberg, 1976). Shaped by this push-pull like organization of the vestibular system, vestibular signals provide key information on neural computation of gravity and body deviation (Angelaki et al., 2004; Merfeld & Zupan, 2002). We tested the impact of increasing loading/unloading on plantar cutaneous information in the mediolateral (ML) axis by comparing the cortical response to plantar sole electric stimulation while a participant (individuals with VH and Controls) stood upright without or with a 1 kg mass added at each wrist.

We hypothesized that adding a weight to both wrists should enhance the detection of body sways through plantar sole mechanoreceptors due to improve sensorimotor integration (assign larger weight to proprioceptive inputs), resulting more accurate encoding of body sways in the gravitational field in the absence of vestibular information. The right posterior parietal cortex (rPPC) is a pivotal region for processing body motion relative the gravitational field (see Pfeiffer et al., 2014, for a review). Observing an increased activity of the rPPC when individuals with VH wore the added mass would support our hypothesis. The approach we used to test this hypothesis was based on the current consensus that functional processing of sensory inputs is associated with the modulation of band-specific neural oscillation power (Haegens et al., 2011; Lebar et al., 2017; Van Ede et al., 2012; Zumer et al., 2014; Pfurtscheller & Lopes da Silva, 1999). We predicted that with the added mass, this modulation would specifically target PPC alpha and beta bands power, which have been associated with the processing of tactile information (Van Ede et al. 2012), notably relative to external frames of reference (Ruzzoli & Soto-Faraco, 2014).

## Participants and methods

### Participants

Ten participants (including 3 women) with bilateral idiopathic vestibular hypofunction took part in the study. Their clinical evaluations are presented in Table 1. They were aged between 44 and 71 (mean 61 years) and their body mass index ranged from 22 and 33 (mean 26, i.e., healthy weight). For all patients, the vestibular pathology started at least 5 years prior to the study. The bilateral vestibular hypofunction was confirmed by a major deficit in the four caloric vestibular tests (the speed of the nystagmus’ slow phase was below 6°/s for all four caloric tests, Barany 1906). The canal function was assessed by measuring the ocular response to low frequency rotations (0.05 Hz, period 20 s), the saccular and the utricular function were tested by measuring vestibular-evoked myogenic potentials (VEMP, respectively on the sternocleidomastoid muscle and on the inferior eyelid). Most patients showed no ocular or muscular response to the rotatory and VEMP tests for utricular function, but they showed muscular response to the VEMP for saccular function. All additional pathologies (e.g., hearing loss) and/or other vestibular pathologies led to the patient’s exclusion from the study. A control group (Controls) was composed of 10 self-declared healthy participants (including 3 women). They were aged between 52 and 64 (mean 58 years) and their body mass index ranged between 22 and 31 (mean 25, healthy weight).

**Table 1:**
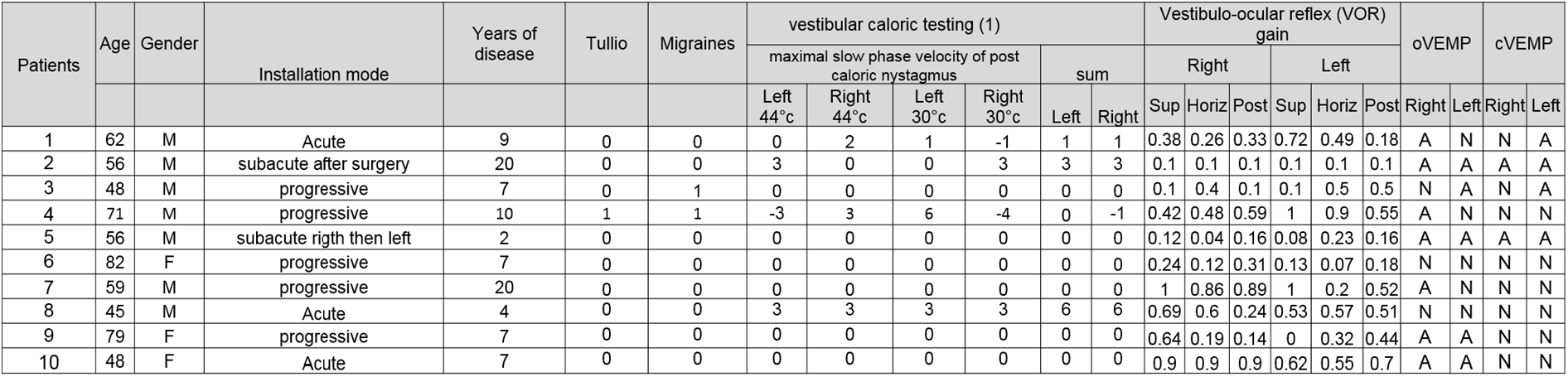
Patients characteristics. Negative values are for the maximal slow phase velocity when nystagmus are beating in opposite side as expected. Ocular (o) and cervical (c) Vestibular Evoked Myogenic Potential (oVEMP) Tests.

All protocols and procedures were conducted in accordance with the ethical standards set out in the Declaration of Helsinki. All participants gave their written informed consent and procedures were approved by the local ethics committee of the Centre d’Explorations Fonctionnelles Oto-neurologiques Falguière (CEFON).

### Task and stimulation procedure

Participants were asked to stand still in a comfortable position with their arms alongside their body and eyes open (Fig.1A). Particular attention was paid to maintain constant the self-selected foot position before each trial to avoid potential effect of stance width on balance control (Day et al., 1993). Before the first session, participants were asked to self-select a comfortable side-by-side foot position and small sticker were positioned in front of each big toe. This allowed us to verify that the feet position remained similar (with an accuracy of a few mm) throughout the experiment.

**Figure 1:**
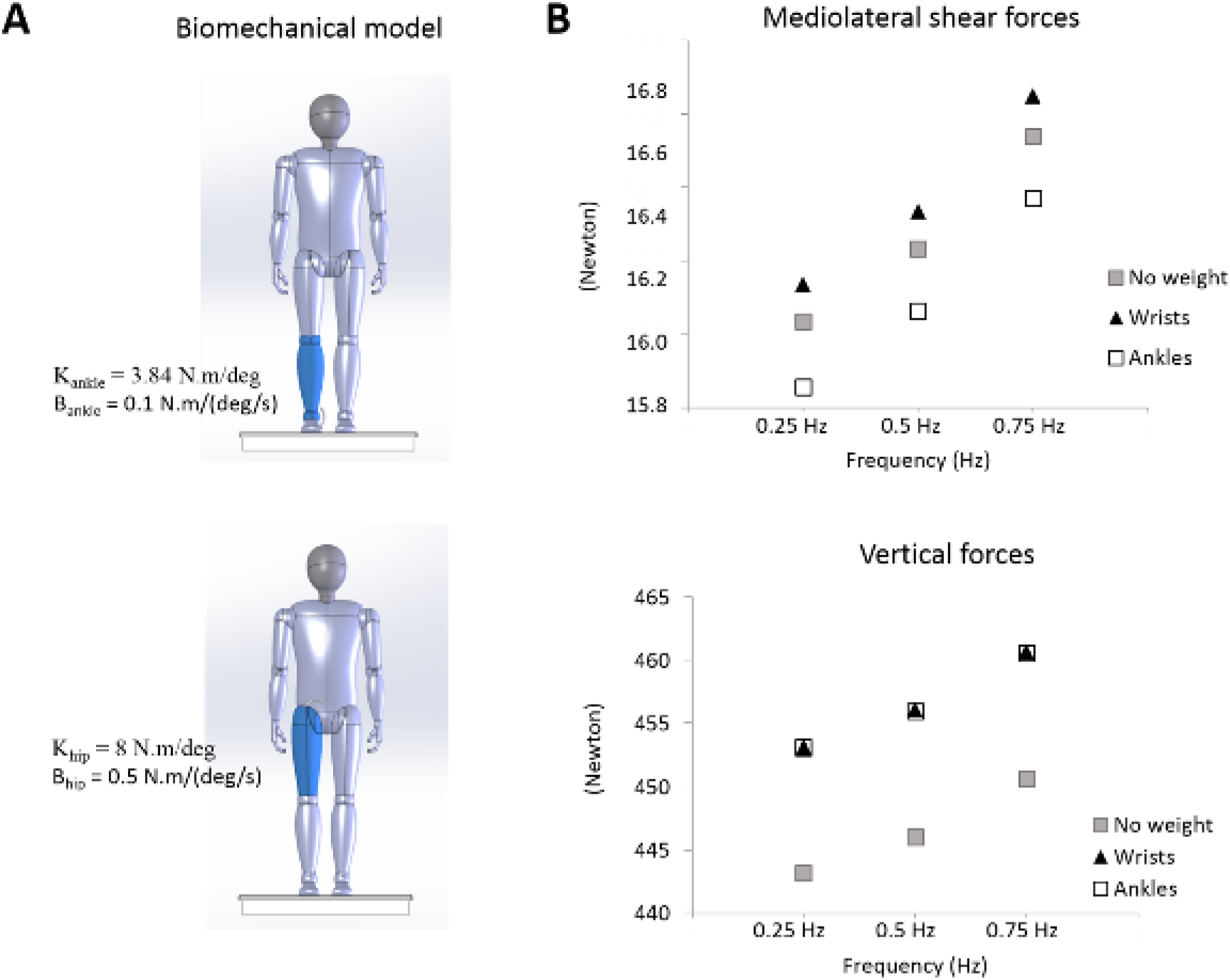
**A**. Biomechanical model. Model represents a double inverted pendulum with movement at the ankle and hip joints. **B** Upper panel depicts peak-to-peak shear forces along the mediolateral axis and the lower panel presents peak-to-peak forces along the vertical axis. Data are for the three center of mass frequencies (i.e., 0.25, 0.5 and 0.75 Hz) and for the three conditions (i.e., without weight, and with the weights at the wrists or at the ankle joints).

Participants were asked to hold their gaze on a small cross fixed on a wall, 3m ahead at eye level. This distance was chosen to avoid vergence eye movements which can be impaired in individuals with VH leading to instability in the mediolateral direction (Kapoula et al., 2013). We recorded cortical activity, and pelvis and head accelerations (see below) of all participants (i.e., VH and Controls) in 16 trials of 15 s. Eight trials were performed while participants wore a 1kg weight bracelet at each wrist (Weight condition) and 8 trials were performed without mass (No weight condition). To avoid potential post-effects of carrying extra mass (e.g., on the frame of reference used for perceiving earth-vertical), the No weight condition (i.e., condition closest to everyday life) was always tested first.

We chose to add 1 kg (i.e., 10 N) at each wrist to increase reaction force under each foot. Such load does not cause disequilibrium while standing upright but influences balance control (e.g., Aruin & Latash, 1995; Mouchnino et al., 2012; Simoneau et al., 2003). Obviously, the added weight increased the body gravitational force. Compared to adding weights at the ankle joints, body sways with weight at both wrists exacerbate the shear forces under each foot (see modeling results below). The current study was conducted in a clinical setting that was not equipped with force platforms. Consequently, we developed a biomechanical model to verify 1) that adding a weight of 20 N altered the vertical and shear reaction forces under both feet during natural body sways and, 2) that this alteration is greater when the weight is added to the wrists (i.e., 10N strapped around each wrist) than to the ankles. For the model, we simulated a double inverted pendulum model (mass = 75 kg, height = 171 cm which correspond to the mean of both groups) with stiffness (K) and damping (B) at the ankle and hip joints (Fig.1A). The biomechanical model only swayed along the frontal plane with an amplitude of 10 mm (i.e., amplitude of center of mass displacement commonly recorded during quiet standing, e.g., Lafond et al., 2004) with frequencies of 0.25, 0.5 or 0.75 Hz. Compared to simulations without the added weight, the outcomes of the model revealed that, as expected, adding a weight of 20 N increased the vertical ground reaction forces under each foot (Fig. 1B) regardless of the added weight location (i.e., wrists or ankles). More importantly, adding weights at the wrists increased the mediolateral shear forces compared to the control condition (i.e., no weight) while adding the weights around the ankle joints decreased these forces (Fig. 1B). Results of the model confirmed that adding weights at both wrists increased the mediolateral shear reaction forces, thereby augmenting the stimulation of the plantar sole mechanoreceptors.

Because of the crossed organization of the ascending sensory pathways, analyzing cortical processes when the laterality of the stimulated foot differs between participants is tricky. In the present study, all participants self-declared being right-handed. We stimulated the left foot for all participants because it is considered as the “postural” foot in most right-handed individuals (Coren, 1993). Electric stimulation of the foot was delivered via a set of 2 electrodes (5 × 9 cm Platinium Foam Electrodes) using an isolated bipolar constant current stimulator (DS5 Digitimer, Welwyn Garden City, UK). Using the same technique as in previous studies (Fabre et al., 2020; Mouchnino et al., 2015), the cathode was located under the metatarsal region and the anode underneath the heel (Fig.2A). Because cutaneous afferents from plantar sole are distributed across the foot sole (with some regional variation, Strzalkowski et al., 2018), the stimulations were deemed to activate most nervous fibres linked to mechanoreceptors. Each stimulation consisted of a single rectangular 10-ms pulse. For each participant, stimulation intensity was set 25% above the tactile detection threshold (i.e., below the motor threshold) and was innocuous. This threshold was estimated before the experimental session using a forced-choice adaptive method (Ehrenstein & Ehrenstein, 1999), while the participants were standing without added mass and were gazing at the fixation cross. The stimulation intensities used for participants with VH and Controls are reported in the Results section. All stimuli were clearly detectable but did not create illusion of body motion, as mechanical stimulation of the foot can do (Kavounoudias et al., 1998). Moreover, a previous study using stimulation intensities similar to those employed here showed that such stimulations are not strong enough to evoke reflex-triggered postural response that could alter normal body sways (Mouchnino et al., 2015).

**Figure 2:**
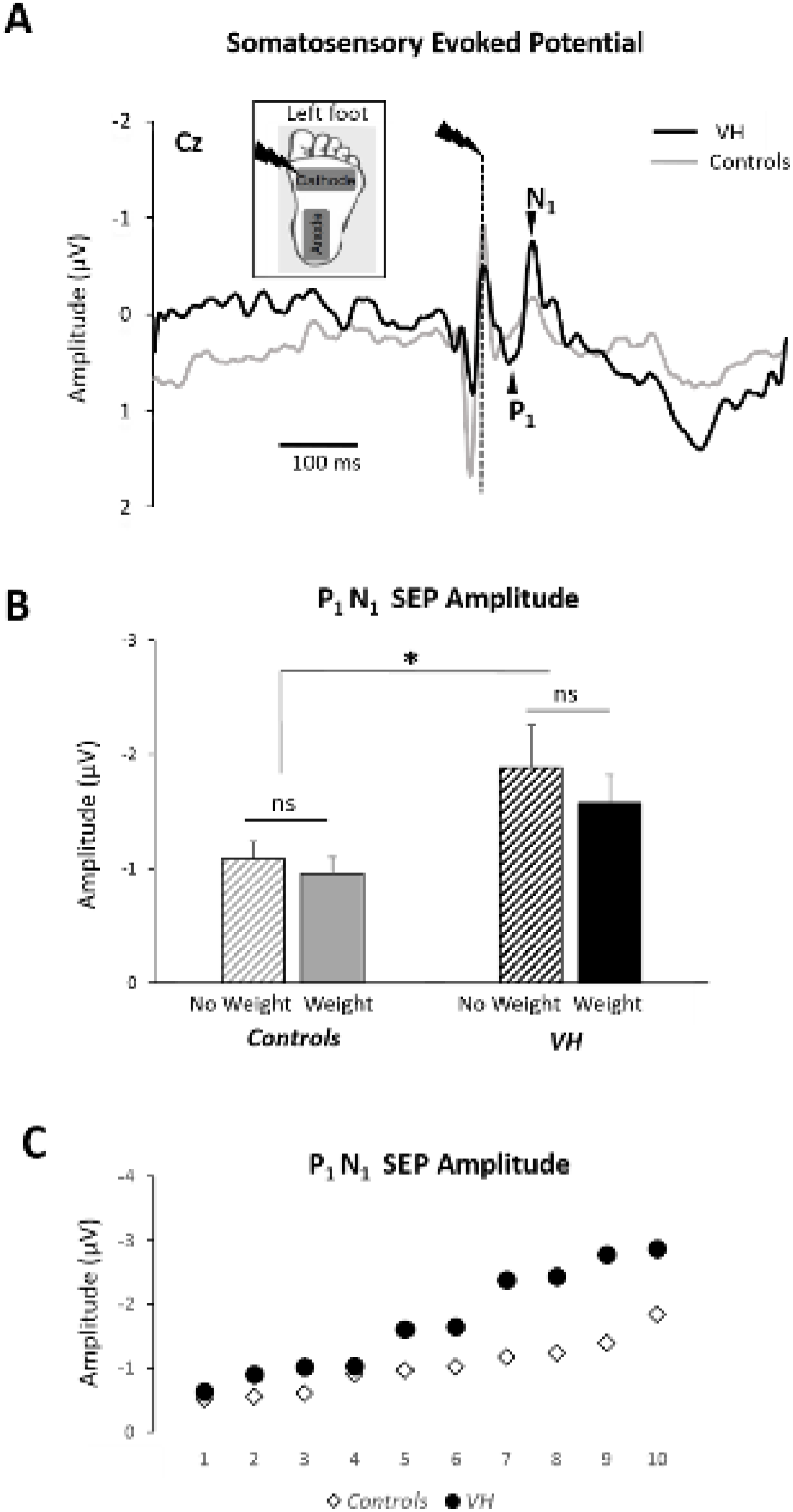
**A**. Position of the stimulation electrodes underneath the left foot (inset). Average of the somatosensory evoked Potential over electrode Cz for a participant in the VH and the Control groups for the No weight condition. The vertical dashed line depicts the onset of the electric stimulation. **B**. Group means amplitude of the P_1_N_1_ SEP evoked by the electric stimulation for the No Weight and Weight conditions (error bars depict standard error of the means). **C**. Averaged Weight and No weight conditions SEP of each participant for all Controls and VH patients. The values are ranked in ascending order for each group.

Fourteen stimulations were delivered in each 15 s trial, giving a total of 98 stimulations for each condition (i.e., No-weight and Weight). The inter-stimulus interval ranged between 800 and 1200 ms to restrain anticipation of the next stimulation by the participants. Because the inter-stimulus intervals were > 800 ms, the evoked response was unlikely influenced by the previous stimulation. When the inter-stimulus intervals was < 500 ms, a decrease in the evoked response was commonly observed due to the previous stimulation (Burke & Gandevia, 1988).

### Electrophysiological recordings and analyses

Electroencephalographic (EEG) activity was recorded continuously from 64 Ag/AgCl surface electrodes embedded in an elastic cap (10-20 system, BioSemi ActiveTwo system: BioSemi, Netherlands). Specific to the BioSemi system, “ground” electrodes were replaced by Common Mode Sense active and Driven Right Leg passive electrodes. The signal was preamplified at the electrode sites and post-amplified with DC amplifiers, digitized at a sampling rate of 1024 Hz (Actiview acquisition program, BioSemi). The signals of each electrode were referenced to the mean signal of all the 64 electrodes. Three external Ag/AgCl electrodes (positioned next to the right and left external canthus and under the left eye orbital) and one from the cap (FP1) controlled for ocular movements and blinks. After artefact rejection based on visual inspection, 87 ± 2% and 83 ± 8% of the trials were included in the analyses for participants with VH and Controls, respectively.

As in previous studies (Duysens et al., 1995, Fabre et al., 2021; Staines et al. 2000), the largest responses to foot stimulations were observed at electrode Cz (vertex). The SEPs were recorded at this electrode which overlays the sensorimotor feet regions located on the inner surface of the longitudinal fissure. These SEPs were obtained by averaging, for each participant and condition (i.e., No-weight and Weight), all synchronized epochs relative to the electrical stimulus (Fig. 2A). We searched, from 25 ms post-stimulus onward (i.e., after stimulation artifact), for the earliest discernible positive and negative peaks. Overall (groups, mass conditions), the latencies of P_1_ and N_1_ were respectively of 59 ± 9 ms and 97 ± 16 ms and were similar as reported in previous studies using similar methods (e.g., Fabre et al., 2020, Duysens et al., 1995). Amplitudes of the P_1_N_1_ were measured peak-to-peak between these positive and negative deflections (Figure 2A).

Time frequency analyses were performed in EEG source space. The neural sources were estimated with dynamical Statistical Parametric Mapping (dSPM, Dale et al., 2000) implemented in the Brainstorm software (Tadel et al., 2011). A three-shell (i.e., scalp, outer skull, and inner skull) sphere boundary element model (BEM) was used to compute the forward model on the anatomical MRI brain template from the Montreal Neurological Institute (MNI Colin27). The data was transformed into time-frequency domain using Morlet wavelet transforms. Due to the Gaussian shape of the frequency response, Morlet wavelets produce smooth-looking time-frequency plots that can be easily interpreted (see Figure 3B, where activities in distinct frequency bands can be identified). We used a 1Hz central frequency (full width at half maximum (FWHM), tc=3sec) that offered a good compromise between temporal and spectral resolutions (Allen & MacKinnon, 2010). We computed alpha (8–12 Hz) and beta (15–25 Hz) event related desynchronization/synchronization (ERD/ERS) using a baseline window (−500 to −50 ms). We purposely selected a 300 ms time window for the ERS/ERD computation [−100; +200 ms relative to the stimulation] because this temporal window allows to include several oscillations cycles. We then extracted the power average of alpha and beta event-related ERS/ERD from 50 to 100 ms (i.e., interval encompassing the P_1_N_1_ SEP). These computations were performed in regions of interest (ROIs) from the right and left posterior parietal cortex (PPC) (see Fig. 3A). These ROIs were defined based on the Destrieux cortical atlas (Destrieux et al., 2010) and had similar numbers of vertices. The right PPC was chosen because it is a key region for integrating tactile information into body representation (Tsakiris et al. 2008, Wang et al. 2016) and on the hypothesis that, for the VH participants, the added mass should enhance the internal representation of body orientation and motion relative to a gravitational reference frame. Consequently, we predicted that the patients’ right PPC should show greater alpha and beta ERDs in the Weight condition than in the No-weight condition. The left PPC, which is not considered to be involved in integrating tactile information into body representation, served as a control ROI. Due to extensive artifact in the beta band, data of 2 participants of the control group were excluded from the time-frequency analysis of the left PPC (i.e., control ROI).

**Figure 3:**
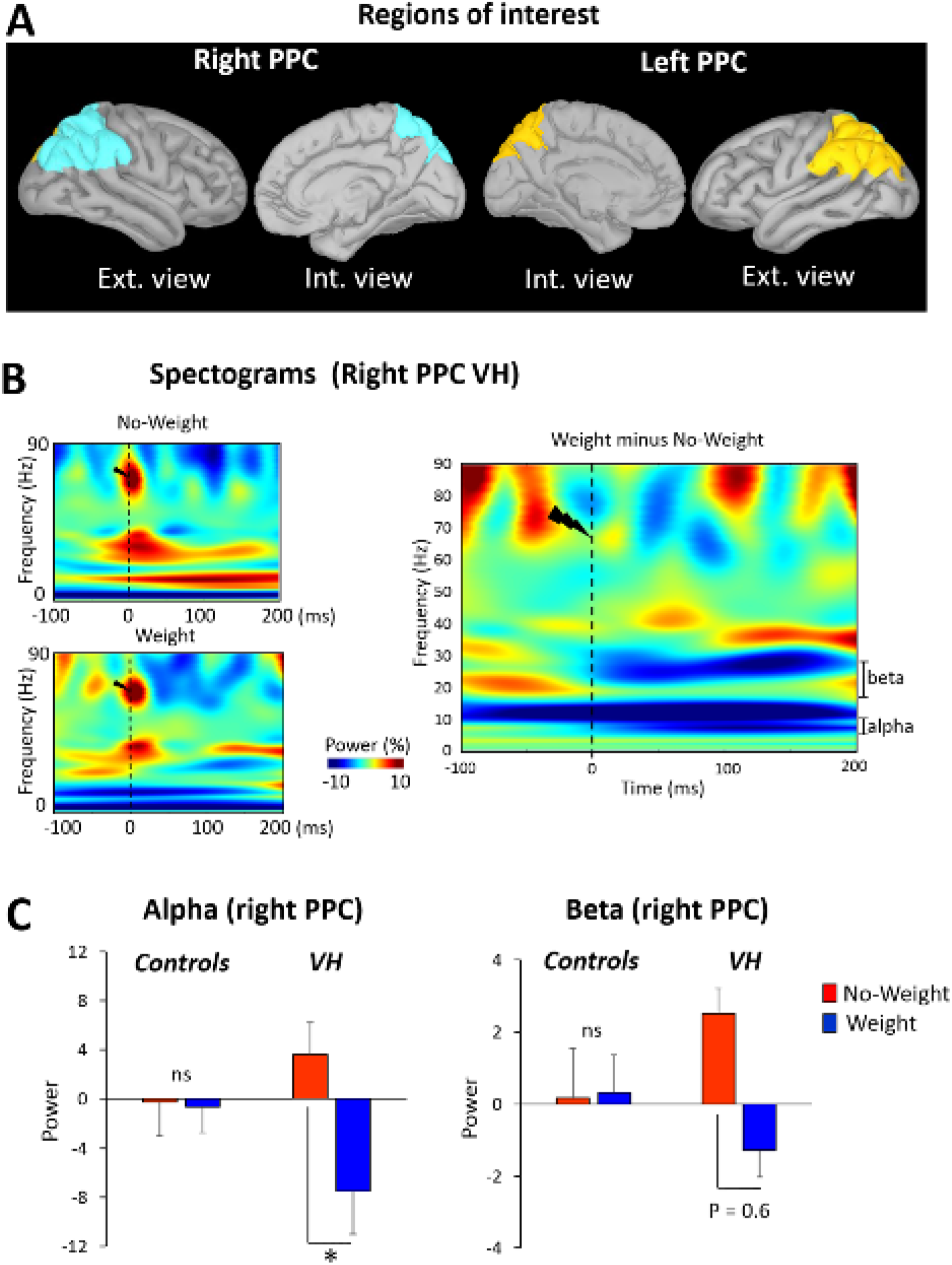
**A**. Location of the regions of interest (ROIs) on the anatomical MRI Colin 27 brain template used to compute cortical activations. Similar ROIs were defined for the left and right Parietal Posterior Cortex (rPPC). **B**. Time-frequency power (ERS/ERD) of the signals by means of a complex Morlet’s wavelet transform applied on the ROIs for each trial of each VH patients then averaged across VH participants in the Weight and No weight conditions (left panel). Cooler colors indicate ERD (event-related desynchronization) while warmer colors indicate ERS (event-related synchronization). Mean difference between Weight and No weight conditions (right panel). Frequency bands up to 90 Hz are illustrated to present changes in brain electrocortical activities for delta, theta, alpha, beta and gamma bands. The artifact caused by the electric stimulation is present at stimulation onset. We epoched the data from −100 to 200 ms to emphasize changes in brain electrocortical activities (ERS/ERD) near plantar sole electrical stimulation. **C**. Group means power for the Alpha (8-12 Hz) and Beta (15-29 Hz) frequency bands computed during [50; 100 ms] time window for the No weight and Weight conditions (error bars depict standard error of the mean).

### Behavioral recordings and analyses

Body stability was assessed by measuring head and pelvis accelerations. Head acceleration was measured with a triaxial accelerometer (Model 4630: Measurement Specialties, USA) placed on the participants’ chin. The accelerometer was securely fixed on a chin-cup. This chin-cup reduced the possibility to move the jaw. Pelvis acceleration was estimated using a triaxial accelerometer (Model ADXL335: Analog Devices, USA) located on the sacrum (i.e., an approximation of the body center of mass, Thirunarayan et al. 1996). Signals from these sensors were recorded at a sampling rate of 1000 Hz and low-pass filtered (Butterworth 4th order, 10 Hz cutoff frequency). Head and pelvis accelerations were analyzed in both anteroposterior (AP) and mediolateral (ML) axes. We rebased, the acceleration timeseries along the AP and ML axes, with respect to their means calculated during the first 500 ms of each 15 s–trial. This procedure allowed a common baseline for all trials. Then, we computed, for each trial, the integrals of the ML and AP time-series to estimate the amount of acceleration in each direction and calculated the mean ML and AP accelerations for each condition and group. These analyses provided an estimate of head and pelvis motion in ML and AP directions for both groups (VH and Controls) and both experimental conditions (i.e., No-Weight and Weight).

### Statistical Analyses

The behavioral and neurophysiological dependent variables were submitted to separate analyses of variance (ANOVA) as the data were normal distributed (Kolmogorov-Smirnov test). For SEP amplitude and latency and alpha/beta ERS/ERD, mixed-design ANOVAs were used for mean comparisons with Condition (No-weight, Weight,) as a within-subject factor and Group (VH, Controls) as a categorical predictor. For head and pelvis accelerations, we used 2 (Group) x (Condition) x 2 (Direction: AP and ML) ANOVAs with repeated measures on the last two factors. For post-hoc analyses, we used Newman-Keuls comparisons. When main effects could be solely explained by a higher order interaction, only the break-down of the interaction are reported. Significance threshold was set at p < 0.05 for all analyses.

## Results

### Foot stimulation perceptual threshold

The mean thresholds for perceiving foot stimulations were 6.95 ± 2.3 mA for the VH and 6.83 ± 1.3 mA for Control groups. The result of a t-test indicated that perceptual thresholds did not differ between both groups (t = −0.23; p = 0.81).

### SEP response to foot tactile stimulation

Consistent with an upregulation of cutaneous inputs for the VH group, the evoked response to the foot cutaneous stimulation was larger in VH than in Control groups (Fig. 2A). This was confirmed by the ANOVA showing that P_1_N_1_ amplitude was greater in VH than in Control groups (Figure 2B-C; F_1, 18_ = 5.84; p = 0.02). Adding a 2 kg mass did not change the SEP amplitude, neither as a main effect (F_1, 18_ = 2.19; p = 0.15) nor as an interaction between Weight x Condition (F_1, 18_ = 0.36; p = 0.55).

For the SEP latency, the ANOVA did not reveal a significant effect of Group (p = 0.82). However, the main effect of Condition failed just short of the conventional 0.05 cut-off value for statistical significance (F_1, 18_ = 4.16; p = 0.07; Weight condition: 58 ms ± 5, No-weight condition: 61 ms ± 8). No Group by Condition interaction was observed (F_1, 18_ = 0.36; p = 0.55).

### EEG spectral analyses

Consistent with an increased activity of the right PPC when adding weight in VH (Fig.3B-C), the alpha power desynchronization increased for the VH group whereas for the Control group, adding weight to the wrists had no significant effect (significant Group by Condition interaction, F_1, 15_ = 6.01; p = 0.02). Similar results were observed for the beta band frequency without reaching the significant value for statistical significance (F_1, 15_ = 4.1; p = 0.062). No significant Condition modulation was observed in the left PPC for both groups, neither for alpha, nor for beta frequency bands (F_1, 15_ = 1.17; p = 0.29 and F_1, 16_ = 3.68; p = 0.072 respectively for alpha and beta).

### Behavioral results

Variables related to balance control were analyzed to compare body sways between VH and Control groups, and to verify whether the added mass altered body sways. For pelvis accelerations (Fig. 4A), the ANOVA revealed a main effect of Group; pelvis accelerations being greater in the Control than in the VH groups (F_1, 17_ = 79; p <0.001). The pelvis acceleration of both groups was greater in the Weight compared to the Non-Weight conditions (main effect of Condition: F_1, 17_ = 8; p = 0.01). The analyses also revealed a significant interaction between Group and Direction (F_1, 17_ = 27; p <0.001). The breakdown of this interaction revealed that pelvis acceleration was greater in the AP direction than in ML for the Control group (p < 0.001), but did not differ between AP and ML directions for the VH group (p = 0.89).

**Figure 4:**
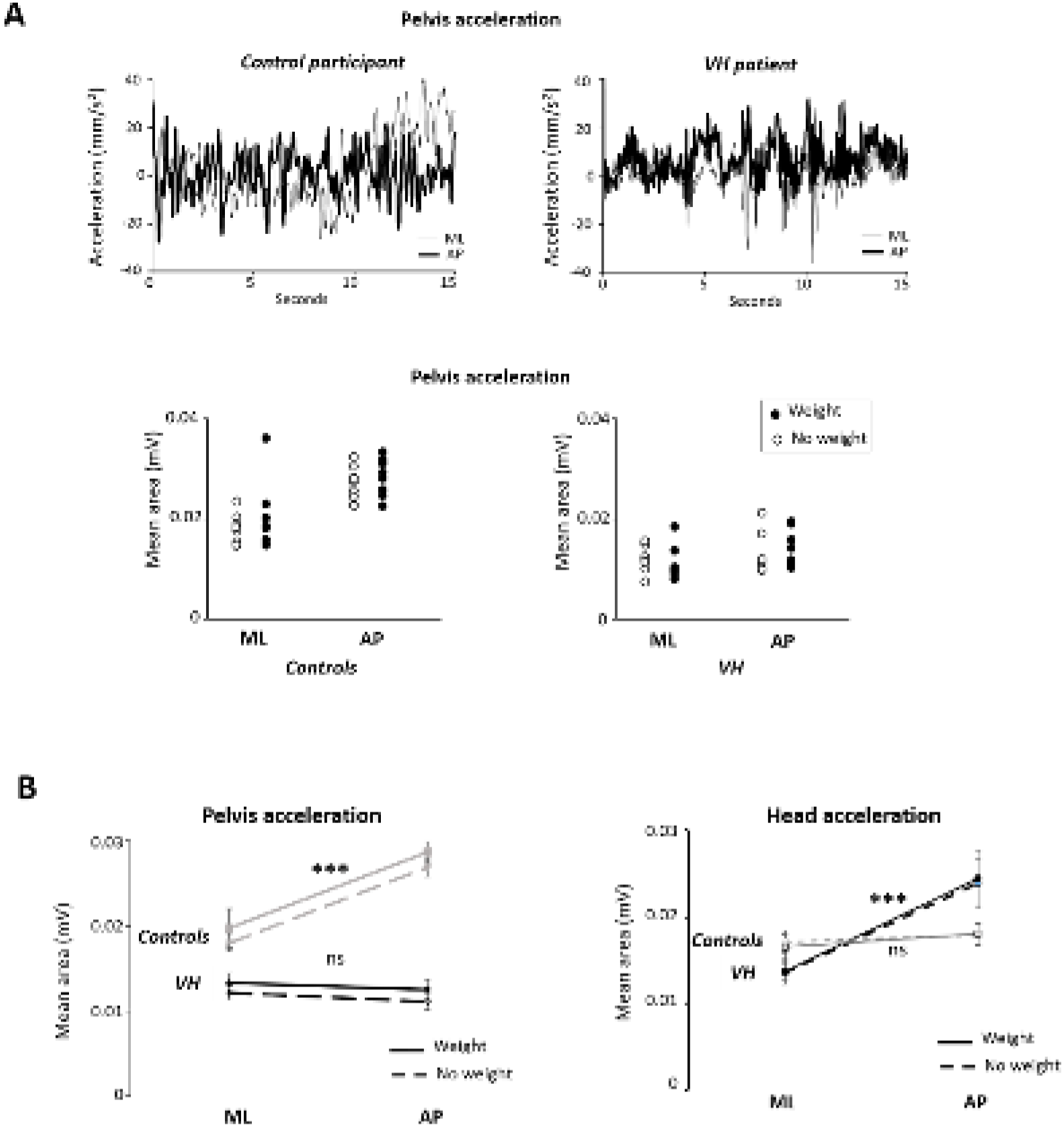
**A**. Upper panel) Pelvis acceleration time-series, along the mediolateral (ML) and antero-posterior (AP) axes, for one representative participant in the Control and VH groups for the No weight condition. Bottom panel) mean of the integration of the pelvis acceleration along the mediolateral (ML) and anteroposterior (AP) directions for each participant for the No weight and the Weight conditions (error bars depict standard error of the mean). **B.** Group means integration of head acceleration along the mediolateral (ML) and anteroposterior (AP) directions for the No weight and Weight conditions (error bars depict standard error of the mean)

Interestingly, for the head accelerations (Fig. 4B), the analyses yielded different results; the changes in head acceleration varied across groups and directions (Group by Direction interaction: F_1, 17_ = 19.2; p <0.001). For the Control group, contrary to what was observed for the pelvis, the head accelerations were similar along the AP and ML directions (p = 0.46) and was uninfluenced by the added mass (p = 0.51). The mean head acceleration of the Control group was not different than the threshold for vestibular detection (0.048 m.s^−2^; Gianna et al. 1996) [*t*-test for means against a value; t_8_ = 1.09, p = 0.30 and t_8_ = 1.03 p = 0.33 respectively for Weight (0.055 ± 0.02 m.^s-2^) and No-weight (0.052 ± 0.01 m.s^−2^) conditions]. Conversely, for VH group, the head accelerations were different between AP and ML directions; head acceleration was greater in AP direction (p < 0.001). No main effect of Group (F_1, 17_ = 0.34; p = 0.56) or Condition (F_1, 17_ = 0.01; p = 0.91) was observed.

## Discussion

Due to the properties of its sensors and their afferents (e.g., otolith neurons) that cannot be turned off (Fernandez & Goldberg, 1976), the vestibular system is a valuable source of information used by the brain to preserve balance during bipedal stance. Being sensitive to linear and angular head accelerations, it provides key information for determining the earth gravity vertical and for estimating how the body deviates from it (Day & Fitzpatrick, 2005). Here, we found evidence that this function could be supported, at least in part, by foot cutaneous inputs in individuals with bilateral vestibular hypofunction. Our results provide valuable insights into the mechanisms underpinning this sensory substitution.

We found that the sensorimotor cortex response to foot cutaneous stimulations was much greater in VH participants than in aged-matched healthy individuals. Although the present experimental protocol does not account for the origin of this finding, the increased VH’s SEP unlikely resulted from an increase in the sensitivity of the plantar sole tactile receptors. Indeed, our results showing similar detection thresholds for foot cutaneous stimulation between VH and Controls provides evidence against this possibility. Distinct potential mechanisms could explain the larger SEP in the VH group. The multimodal cells, where vestibular and somatosensory inputs converge, could have augmented their responsiveness to cutaneous stimulations following reduced vestibular inputs. Such cells are present, for example, in the somatosensory cortex (Zarzecki et al., 1983; Guldin and Grüsser, 1998) and in the thalamus (Lopez and Blanke, 2011). Such trade-off mechanism between somatosensory and vestibular responsiveness has been observed in upright standing healthy individuals by Mian and Day (2014). These authors showed that the gain of the vestibular-evoked balance response was lessened during finger tactile stimulation most likely enhancing the weight of somatosensory inputs as shown by Bolton et al. (2011) in participants touching lightly an external support. Besides, the thalamus, which is the principal sensory relay station, can also send potentiated inputs to the brain through the effect of the reticular nucleus which intercepts and modulates all corticothalamic (including from prefrontal cortex) and thalamocortical communications (Guillery et al., 1998; Zikopoulos and Barbas, 2006). Because cells that respond exclusively to vestibular stimulations have not yet been discovered in the brain, it is unlikely that greater SEP in participants with VH resulted from cross-modal plasticity. This phenomenon allows for instance cells which respond to visual stimulation in sighted individuals to be sensitive to cutaneous inputs in the blinds (Bavelier and Neville, 2002; Sathian and Stilla, 2010).

Our behavioral data support the suggestion that somatosensory compensation occurred in absence of vestibular cues. For the VH group, pelvis accelerations were reduced along both directions (i.e., AP and ML) while for the Control group, we observed a reduction in head accelerations in both directions (see Fig. 3). This reduced head accelerations could imply that in healthy individuals, the head served as a reference frame for controlling upright standing and locomotion. This strategy requires vestibular functioning (Bent et al., 2004; Horak et al., 2002; Pozzo et al., 1991). Note that, along the frontal plan, the mean head acceleration of the control participants was not different than the acceleration threshold for vestibular detection (Gianna et al., 1996). Functionally, such near-vestibular threshold head acceleration puts the vestibular system in an optimal state for being stimulated in case of unexpected loss of balance.

For the participants with VH, we observed smaller pelvis accelerations. The brain, in the absence of vestibular information, likely determined body sways amplitude and direction with respect to the base of support through somatosensory information (e.g., lower limb somatosensory and plantar sole afferents signals). This change in reference frame involves a tight control of the pelvis. In particular, the body sways can be regulated within a small area by reducing the amplitude of the centre of pressure displacement (not recorded in the present study) and increasing its frequency. Such tactile exploration, by changing the pressure onto the ground with the feet, likely contributes to improve the signal-to-noise ratio of sensory cues and to determine the limit of stability (Fabre et al. 2021; Latash et al., 2003; Murnaghan et al., 2011). The increase in plantar sole afferents through this strategy could be particularly important in the absence of vestibular information. Larger upregulation of cutaneous inputs could have contributed to the greater SEP observed in the participants with VH compared to controls. Attentional processes, which are known to increase the excitability to sensory input (Eimer & Forster, 2003; Sambo & Forster, 2011; Bolton & Staines, 2012) could be part of the mechanisms that contributed to this sensory upregulation.

Besides and, perhaps more surprisingly, for participants with VH, we did not observe difference in the pelvis accelerations along the AP and ML directions. In contrast, the healthy individuals exhibited larger pelvis accelerations in the AP compared to the ML directions as reported in previous studies (Collins & de Luca, 1993; Lord et al., 1999; Johnson-Hilliard et al., 2008; Winter et al., 1996). Less accurate estimation of body sways amplitude and direction, due to the absence of vestibular information, likely increases the perception of instability leading to tight control of body sways along the frontal and sagittal planes. A tight control of body sways is commonly observed in healthy individuals when they are standing on an elevated surface (Adkin et al., 2000; Carpenter et al., 1999; Lim et al., 2017) or in patients with postural phobic vertigo (Krafczyk et al., 1999). The increased risk of injury leads to tight balance control.

Our second hypothesis was that potentiating the differential pressure between both feet when standing could contribute to restoring the internal representation of the patients’ body orientation and motion relative to a gravitational reference frame hosted in the right PPC (Pérennou et al., 2008; Dai et al., 2021a, b; Pfeiffer et al., 2014). In the present study, this potentiation was done by adding a 1 kg mass to both wrists. Our biomechanical model indicated that this procedure magnified the differential pressure between both feet, and more specifically the shear forces. The analyses of the alpha and beta power recorded in VH participants are consistent with an enhanced internal presentation of the body when standing with the added mass. We found that adding a small weight on each hanging wrist resulted in a decrease of the right PPC alpha and beta power in the VH but not in the Control groups. The powers of the alpha and beta oscillations are inversely related to cortical excitability and to the reliance on sensory processing, respectively (Neuper & Pfurtscheller, 2001; Romei et al., 2008; Sauseng et al., 2009). For VH participants, the decreased alpha and beta power observed in the right PPC might entail integration of cutaneous inputs into the representation of body orientation relative to the gravitational field. This cortical region is known to receive thalamic projections of cutaneous cues through either direct (Padberg et al. 2009; Impieri et al. 2018) or indirect (i.e., through SII/parietal ventral, Cavada and Goldman-Rakic 1989; Disbrow et al. 2003) pathways. In VH participants, the contribution of the PCC to balance control could have therefore been enhanced by exteroceptive tactile afferents in the Weight condition. This would be consistent with findings suggesting that the reliance on internal representations for controlling balance increases in contexts with impaired sensory inputs (Blouin et al., 2007; Fabre et al., 2020). The importance of cutaneous inputs in building a representation of the body in space has also been suggested by observations made by Day & Cole (2002) in a study with a deafferented patient (IW). The patient, who suffered from a rare lack of cutaneous and proprioceptive inputs (see Cole & Waterman, 1995), experienced a “floating sensation” and a “loss of contact with the world” when receiving galvanic vestibular stimulations (GVS) while seated. This unusual effect suggests an inability to create an accurate representation of the body in space in the absence of cutaneous inputs, even when vestibular information is preserved.

In conclusion, we found that plantar cutaneous inputs are upregulated in individuals with major vestibular information deficits. Such sensory upregulation points to an increased importance of cutaneous input in balance control in these patients. The contribution of foot tactile information in VH was also suggested by their change in balance control and postural organization, which were based on reduced pelvis acceleration rather than head acceleration. Finally, increasing loading/unloading under both feet with small weights, we were able to identify within the right PPC altered somatosensory dynamics (i.e., increased alpha-beta desynchronization). This supports our hypothesis of an increased processing of tactile information relative to an external frame of reference to enable building up an internal representation of body state.

## Acknowledgements

We are thankful to the Association Française de vestibulopathie bilatérale idiopathique and the patients for their participation to the study. We thank F. Buloup for developing the software Docometre used for data processing and acquisition. The authors are grateful to Alec Blouin and Guillaume Vinay for their technical assistance. They also thank Elsa Baroghel for reviewing the English.

